# Targeting CHI3L1 in Alzheimer’s Disease: Optimization of G721-0282 and Functional Evaluation in Astrocyte Models

**DOI:** 10.64898/2026.01.13.699206

**Authors:** Baljit Kaur, Hossam Nada, Longfei Zhang, Moustafa Gabr

**Author notes:** To whom correspondence should be addressed: Moustafa T. Gabr.

## Abstract

Alzheimer’s disease (AD) involves astrocytic dysfunction characterized by impaired lysosomal activity, defective amyloid clearance, and neuroinflammation, processes strongly regulated by the inflammatory effector CHI3L1. **G721-0282**, a reported CHI3L1-binding small molecule with demonstrated modulation of downstream signaling pathways including MAPK and STAT3, provides a validated chemical starting point for targeting CHI3L1-driven astrocytic pathology in AD, but exhibits suboptimal potency and drug-like properties that limit its translational potential. We therefore performed a virtual screening of commercially available analogues of **G721-0282** to enable structure-guided optimization, generating a detailed structure–activity map and prioritizing 24 derivatives. Biophysical analyses identified compound **G721-0377** as the most promising candidate, with optimized substitutions improving CHI3L1 binding affinity. Compound **G721-0377** also exhibited favorable pharmacokinetic properties, including improved solubility, balanced permeability, reduced microsomal clearance, and an enhanced cardiac safety margin. Functionally, **G721-0377** uniquely reversed CHI3L1-induced astrocytic dysfunction, restoring amyloid uptake, lysosomal proteolysis and acidification, suppressing CHI3L1 and IL-6 secretion, and inhibiting NF-κB activation to levels comparable to a neutralizing anti-CHI3L1 antibody. In contrast, compounds **G721-0179** and **G857-1069** showed minimal activity. Collectively, these findings establish **G721-0377** as a next-generation CHI3L1 inhibitor with improved affinity, safety, and robust functional efficacy, supporting its further development as a disease-modifying therapeutic for AD.

## Introduction

Alzheimer’s disease (AD) is a multifaceted neurodegenerative disorder that impacts millions globally and currently lacks effective disease-modifying treatments.^1,2^ Although amyloid-β (Aβ) plaques^3^ and tau neurofibrillary tangles^4^ define the core neuropathology, these aggregates develop within a broader context of glial dysfunction,^5^ impaired proteostasis,^6^ and chronic neuroinflammation,^7,8^ which significantly contribute to disease progression. Among glial cell types, astrocytes have gained particular attention as active regulators of neuronal homeostasis^9^ and as critical contributors to disease-associated pathology.^10,11^

Astrocytes play essential roles in maintaining brain health, including regulation of synaptic activity,^12^ metabolic support of neurons,^13^ and clearance of extracellular Aβ through receptor-mediated endocytosis and lysosomal degradation.^14^ In AD, astrocytes undergo maladaptive reactive changes marked by impaired Aβ uptake, disrupted lysosomal function, and heightened secretion of inflammatory mediators.^15^ These alterations compromise their neuroprotective roles and contribute to the accumulation of toxic protein species.^16^

Chitinase-3-like 1 (CHI3L1), commonly referred to as YKL-40, has gained attention as a key astrocyte-related factor in Alzheimer’s disease (AD).^17,18^ Elevated levels of CHI3L1 have been consistently reported in the cerebrospinal fluid and brain tissue of individuals with AD, and higher concentrations often correspond with more severe cognitive decline and neuropathological burden. ^19,20^ Within the brain, CHI3L1 is predominantly produced by reactive astrocytes,^21,22^ and its expression is frequently observed in areas enriched with amyloid plaques or vascular amyloid deposition, indicating a close association with local amyloid-related stress.^23,24^ Clinically, increased CSF (Cerebrospinal Fluid) CHI3L1 aligns with alterations in established AD biomarkers, including Aβ_42_ and phosphorylated tau, and can help identify individuals at risk of progressing from presymptomatic to symptomatic stages of the disease.^25^

Although CHI3L1 lacks enzymatic activity, it participates in signaling pathways that influence inflammation, extracellular matrix dynamics, and cellular stress responses.^26–29^ Experimental studies show that manipulating CHI3L1 levels can alter amyloid accumulation and modify astrocytic reactivity, suggesting that CHI3L1 plays a regulatory role in how astrocytes respond to chronic protein aggregation.^30,31^

Recent functional studies extend this concept by demonstrating that CHI3L1 directly disrupts astrocytic Aβ handling.^32,33^ Human genetic analyses show that reduced CHI3L1 expression is associated with slower AD progression, while mouse models indicate that CHI3L1 deletion enhances astrocyte and microglial phagocytosis, reduces amyloid plaque burden, and promotes lysosomal activation.^34^ These findings support a functional role for CHI3L1 in suppressing glial clearance mechanisms and linking astrocytic activity to disease progression. Overall, current evidence supports CHI3L1 as both a promising biomarker and a modulator of astrocyte-driven pathology in Alzheimer’s disease.

Despite its central role, CHI3L1 has largely remained “undruggable,” with few small molecules reported to bind or inhibit its function.^35,36^ Recent advances in sensitive biophysical screening and structure-based drug discovery now offer tools to overcome this challenge,^37,38^ although drug delivery remains constrained by factors such as blood-brain barrier permeability.

Among reported modulators, **G721-0282** has been identified as a CHI3L1-binding small molecule with moderate activity in previous studies, where it influenced signaling pathways such as MAPK and STAT3.^39–42^ However, the native scaffold exhibits limited potency and variable target engagement, with moderate binding observed in biophysical assays and inconsistent functional outcomes across different models. These findings indicate that scaffold optimization is needed to improve binding affinity, enhance inhibition of CHI3L1-mediated processes, and achieve consistent biological effects. Its activity in Alzheimer’s disease-relevant contexts had not been explored. We selected **G721-0282** as a starting scaffold due to its confirmed CHI3L1 engagement and aimed first to enhance its binding affinity with CHI3L1. Following optimization, we evaluated the compound’s functional impact in AD-relevant assays, including astrocyte-mediated Aβ uptake, lysosomal function, and NF-κB-dependent inflammatory signaling, thereby systematically improving CHI3L1 engagement while assessing its mechanistic relevance in AD.

## Results and Discussion

### Chemical Features of Virtual Screening Hits Identified by Molecular Docking

To optimize CHI3L1 modulation for Alzheimer’s disease-relevant astrocytic dysfunction, we conducted a virtual screening campaign of commercially available **G721-0282** analogues. This approach aimed to identify derivatives predicted to exhibit improved binding affinity and favorable interactions with key CHI3L1 residues, allowing prioritization of compounds for biophysical and functional evaluation. Emphasis was placed on both docking performance (Table S1) and structural diversity to ensure broad coverage of chemical space, resulting in **24** structurally diverse derivatives selected for further investigation (Figure 1). Descriptive analysis of these hits revealed substantial chemical diversity while retaining a common core scaffold. The compounds included aliphatic substituents, monocyclic and fused aromatic systems, and heteroaromatic rings, as well as small ether, ester, and amide functionalities. Aromatic groups with varying electronic characteristics were represented, along with fused frameworks such as naphthalene- and benzodioxole-type structures. In addition, variability was observed in linker motifs, including linear and branched alkyl chains and ether-containing linkages, reflecting differences in steric bulk and polarity across the hit set.

**Figure 1.**
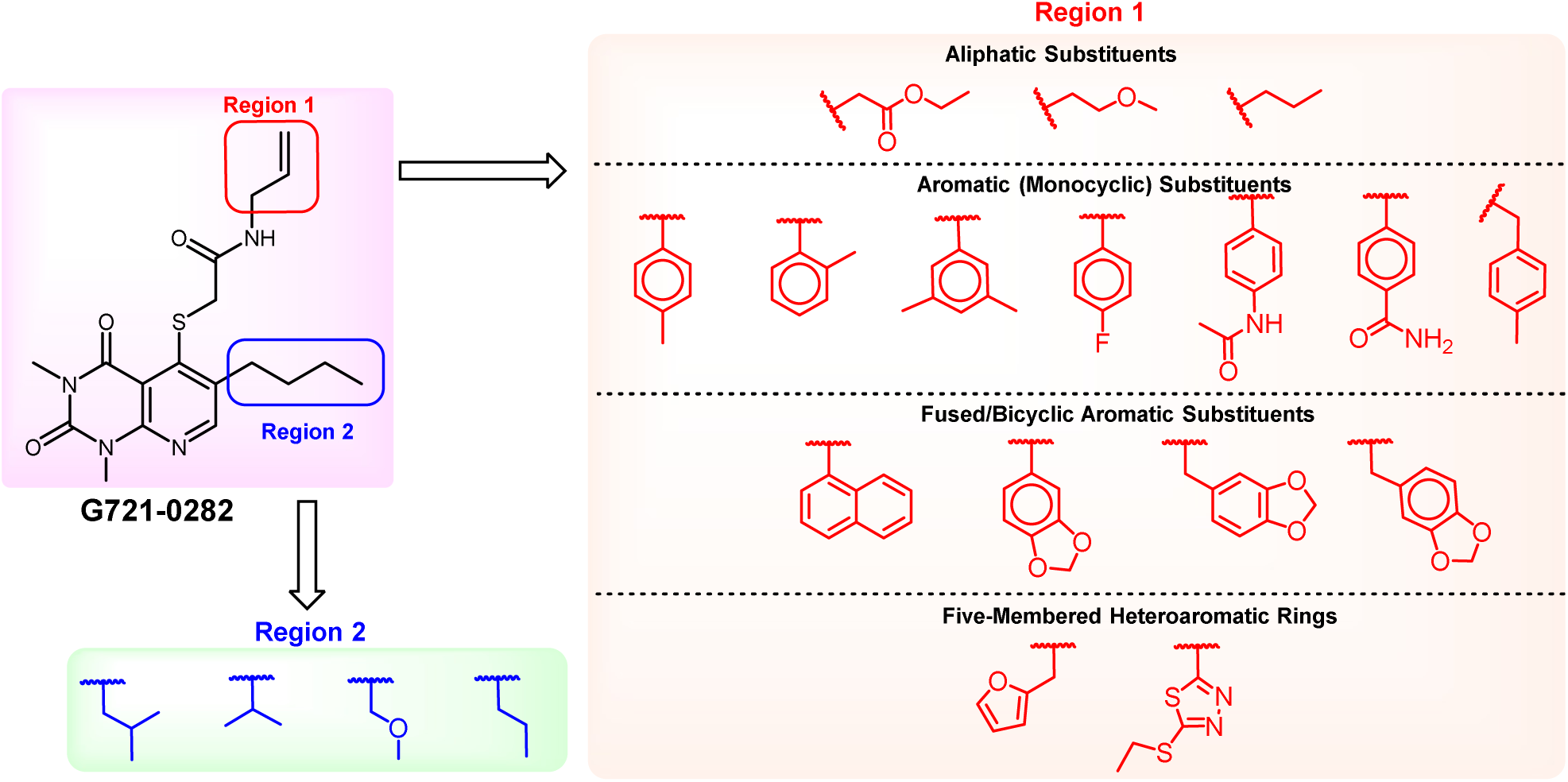
Comparison of Region 1 and Region 2 moieties between the parent compound **G721-0282** and 24 virtual screening hits.

### Biophysical Screening

Before screening the compound library, we first assessed the binding of the parent compound **G721-0282** with CHI3L1 protein (Figure 2). A binding check on Dianthus confirmed that **G721-0282** binds to the CHI3L1 protein, showing a signal-to-noise ratio of 37.26 and a signal of 15.36 (Figure 2B). However, as we reported in our previous publication,^36^ **G721-0282** failed to produce a dose-response curve using the TRIC mode of the Monolith instrument. In the current study, we employed the more sensitive Monolith X instrument, which uses spectral shift detection. This platform successfully generated a dose-response curve for **G721-0282**, for which the TRIC trace had previously been ineffective. However, the binding of **G721-0282** to CHI3L1 remained very weak, with a K_D_ of 155 uM (Figure 2C).

**Figure. 2.**
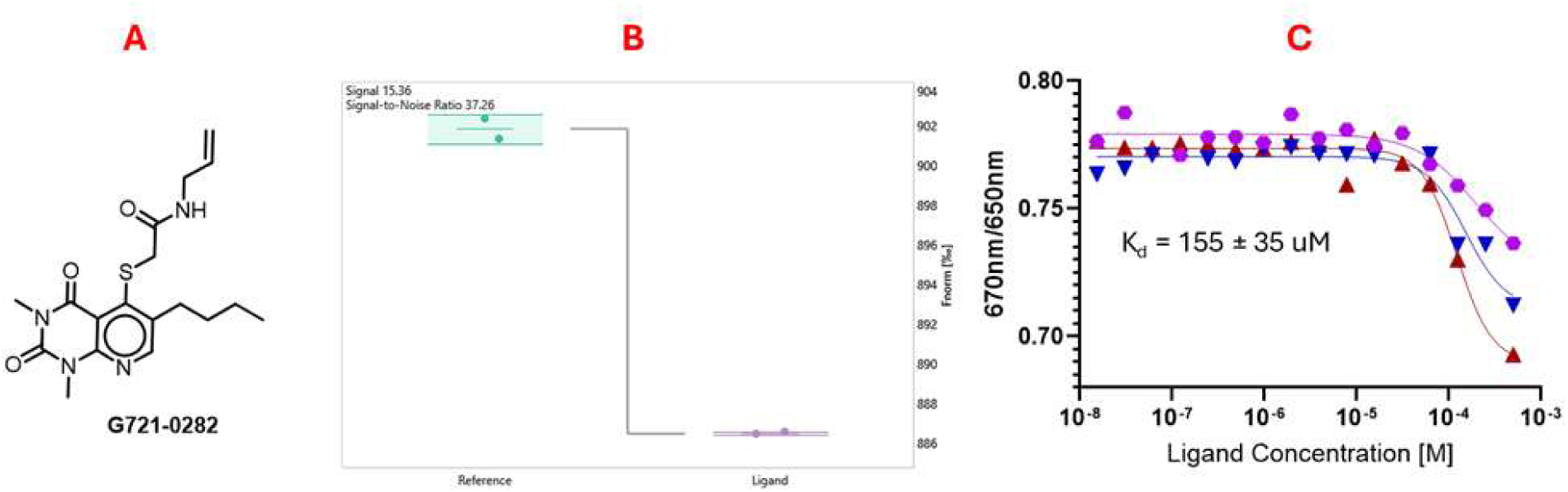
(**A**) Chemical Structure of **G721-0282**; (**B**) Binding check of **G721-0282** on dianthus; (**C**) Dose response **of G721-0282** using spectral shift technique on monolith X.

The selected top 24 compounds were procured from ChemDiv, and stock solutions were prepared in DMSO. Initial screening was conducted using the MST-based Dianthus instrument. Following the same procedure reported previously,^43^ preliminary binding assessment was performed at a compound concentration of 250 µM with a final DMSO concentration of 2.5%. His-tagged human CHI3L1 protein was labeled with the NTA second-generation dye (Nanotemper) in a 2:1 ratio and incubated with the compounds at a 1:1 ratio. Screening was performed on the Dianthus platform, and all experiments were carried out in triplicate. Out of the 24 compounds screened, 10 were identified as initial hits (Figure 3A). Comparison of the initial fluorescence values of the compounds with the DMSO control suggested that four of these could be false positives due to autofluorescence or quenching effects (Figure 3B). To verify this, control experiments were performed (Figures S1 & S2). None of the compounds exhibited significant autofluorescence or quenching.

**Figure 3.**
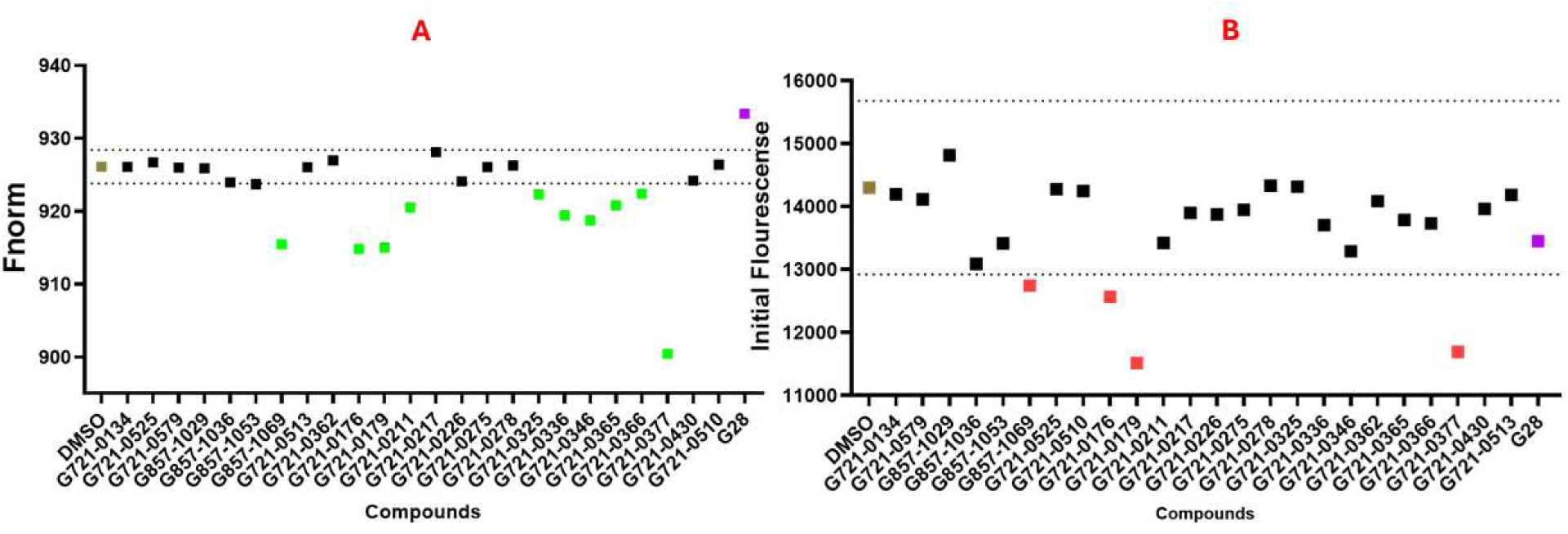
(**A**) Primary screening of compounds at 250 μM (2.5% DMSO). Green, brown, black, and purple represent potential hits, the negative control/reference (buffer with 2.5% DMSO), non-binders, and the positive control (**G28**),^36^ respectively. (**B**) Comparison of the initial fluorescence of the compounds with the DMSO-containing reference; red highlights compounds with a likelihood of MST interference due to autofluorescence or quenching.

After the initial screening, all ten hits were evaluated using dose-dependent MST analysis on the Monolith instrument. Among the ten compounds tested, only three compounds – **G721-0377**, **G721-0179**, **and G857-1069** showed clear dose-response curves with CHI3L1, indicating their potential as genuine binders (**Figure 4**). These results suggest that replacing the allyl group with a *p*-tolyl group in **G721-0179** enhances its binding affinity for CHI3L1. Interestingly, substituting the parent compound’s four-carbon alkyl chain with an isobutyl group, along with replacing the allyl moiety with a benzamide group, further improves binding affinity as reflected by the K_d_ shift from 155 µM in the parent compound to 45 µM in **G721-0377**. In contrast, replacing the four-carbon chain with a three-carbon chain in **G857-1069** reduces binding affinity, as shown by the increased K_d_ of 236 µM (**Figure 4**). Out of these three hits, we selected **G721-0377** and **G721-0179** for further studies along with **G721-0282**.

**Figure 4.**
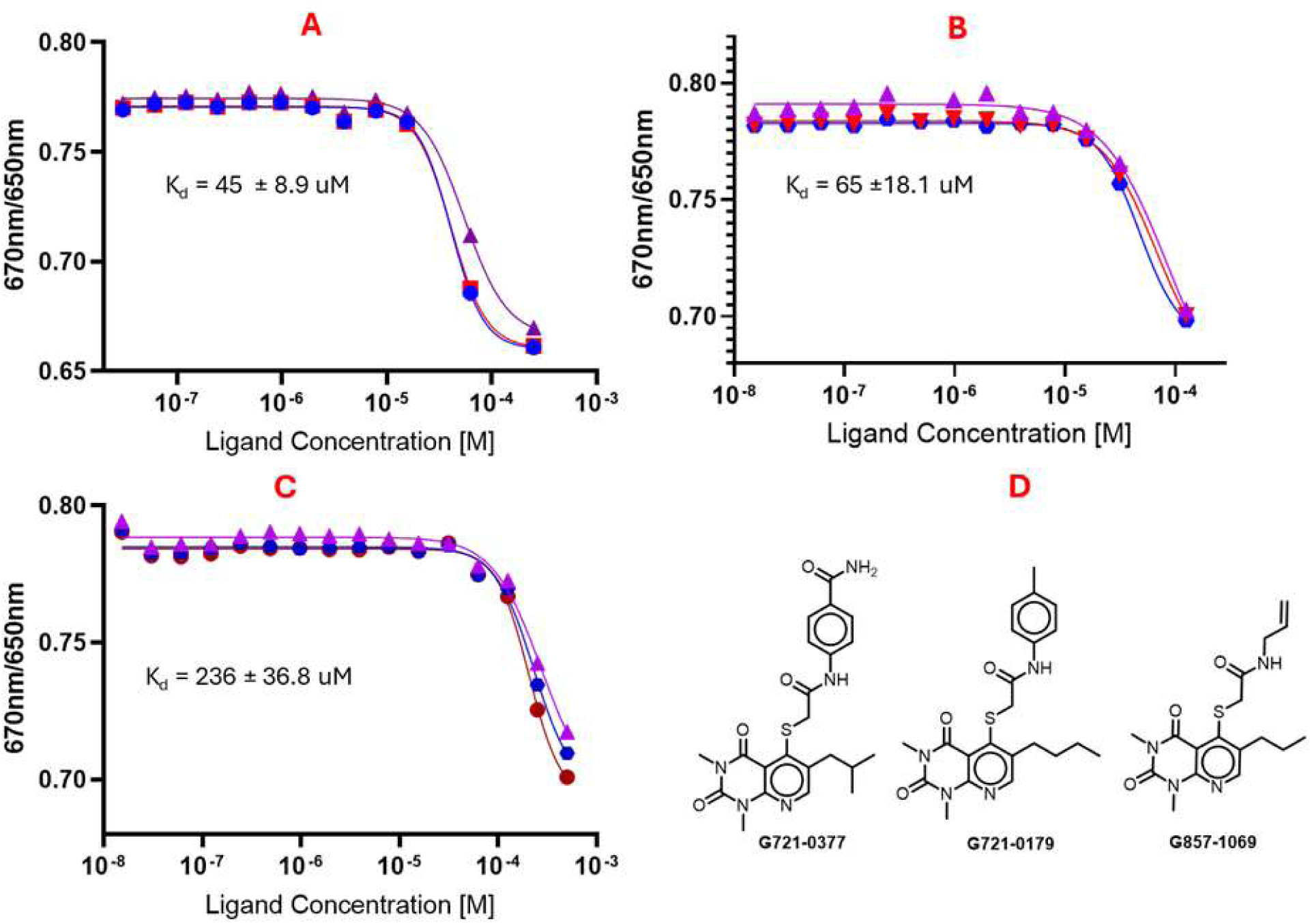
Dose response curves of compounds interacting with hCHI3L1: (A) **G721-0377**, (B) **G721-0179**, and (C) **G857-1069** and (D) Chemical structures of hits.

### PK Profiling

The PK properties of small molecule drug candidates for AD impose particularly stringent requirements, as successful candidates must combine sufficient central nervous system (CNS) exposure with metabolic stability, favorable safety margins, and appropriate physicochemical balance to support chronic dosing. Within this context, the in vitro PK profile of the **G721** series provides key insight into the relative developability of these compounds as potential AD therapeutics.

**G721-0282**, the original lead from this series, exhibited moderate lipophilicity (logD_7.4_ = 3.2) and acceptable passive permeability; however, it suffered from rapid clearance in both mouse and human systems, reflected by short plasma half-lives (<1 h) and high intrinsic clearance in liver microsomes (Table 1). Such a profile is generally incompatible with sustained CNS exposure and would be expected to limit overall brain exposure in vivo unless compensated by frequent or high dosing, which is undesirable in the context of a chronic neurodegenerative disease. In addition, the hERG liability observed for **G721-0282** (IC_50_ = 4.8 µM) raises concerns regarding cardiac safety, further constraining its translational potential. **G721-0179**, which incorporates a more lipophilic aryl carbamate substituent, demonstrated substantially enhanced passive permeability (PAMPA Papp = 5.3 × 10^-6^ cm/s), consistent with its increased lipophilicity (logD_7.4_ = 4.4). While elevated lipophilicity is often beneficial for BBB penetration, this gain came at the cost of markedly reduced aqueous solubility (5.1 µM) and extremely high plasma protein binding (99.4%), both of which are unfavorable for achieving sufficient free drug concentrations in the CNS (Table 1). Moreover, this compound retained high intrinsic clearance similar to **G721-0282**, indicating that the increased lipophilicity did not translate into improved metabolic stability. Importantly, **G721-0179** showed the strongest hERG liability among the series (IC_50_ = 3.1 µM), substantially narrowing its therapeutic window and rendering it unsuitable for chronic administration in a frail elderly population. In contrast, **G721-0377** exhibited the most balanced PK and safety profile across the series and addresses many of the limitations observed for the other antagonists (Table 1). In quantitative terms, **G721-0377** achieved significantly improved solubility (32.7 µM) while maintaining favorable permeability (PAMPA Papp = 3.8 × 10⁻⁶ cm/s), reflecting a more optimal balance between polarity and hydrophobicity (logD_7.4_ = 3.8). This balance translated directly into improved metabolic stability, as evidenced by lower intrinsic clearance in both mouse and human microsomes (24 and 22 mL/min/mg, respectively) and correspondingly extended plasma half-lives in both species. From the perspective of CNS pharmacokinetics, such improvements are highly desirable, as reduced clearance is one of the most critical determinants of sustained brain exposure.

**Table 1.**
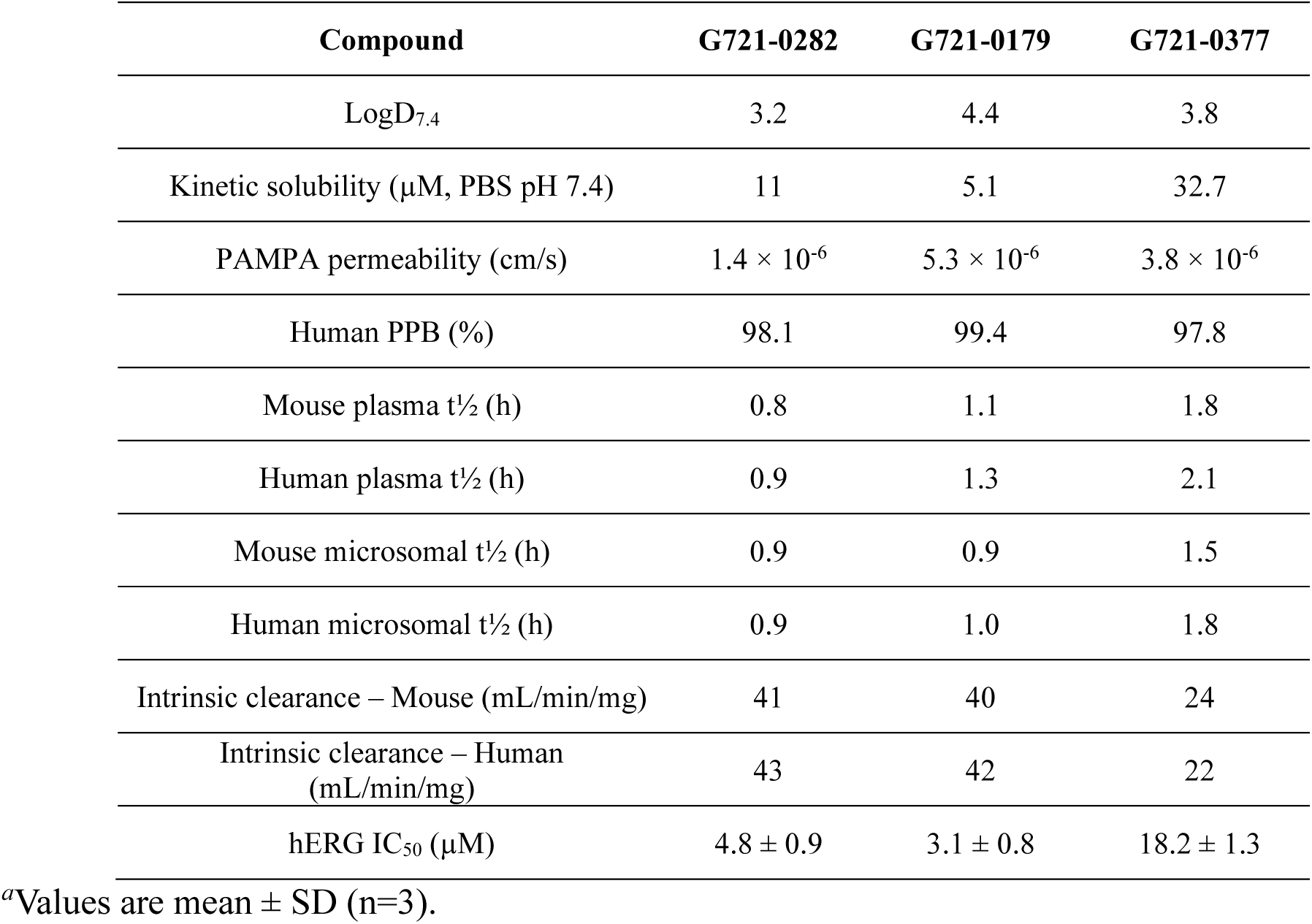
Assessment of the in vitro PK properties of G721-0282, G721-0179, and G721-0377.^a^.

| Compound | G721-0282 | G721-0179 | G721-0377 |
| --- | --- | --- | --- |
| LogD<sub>7.4</sub> | 3.2 | 4.4 | 3.8 |
| Kinetic solubility (μM, PBS pH 7.4) | 11 | 5.1 | 32.7 |
| PAMPA permeability (cm/s) | 1.4 × 10<sup>-6</sup> | 5.3 × 10<sup>-6</sup> | 3.8 × 10<sup>-6</sup> |
| Human PPB (%) | 98.1 | 99.4 | 97.8 |
| Mouse plasma t<sub>1/2</sub> (h) | 0.8 | 1.1 | 1.8 |
| Human plasma t<sub>1/2</sub> (h) | 0.9 | 1.3 | 2.1 |
| Mouse microsomal t<sub>1/2</sub> (h) | 0.9 | 0.9 | 1.5 |
| Human microsomal t<sub>1/2</sub> (h) | 0.9 | 1.0 | 1.8 |
| Intrinsic clearance – Mouse (mL/min/mg) | 41 | 40 | 24 |
| Intrinsic clearance – Human (mL/min/mg) | 43 | 42 | 22 |
| hERG IC<sub>50</sub> (μM) | 4.8 ± 0.9 | 3.1 ± 0.8 | 18.2 ± 1.3 |

<sup>a</sup>Values are mean ± SD (n=3).

Notably, **G721-0377** also demonstrated a substantially improved cardiac safety profile, with an hERG IC_50_ of 18.2 µM, representing an approximate six-fold safety margin enhancement relative to **G721-0282** and **G721-0179** (Table 1). This improvement likely arises from the increased polarity and altered electronic properties introduced by the para-amide substituent, which may reduce nonspecific ion channel interactions. Given that cardiovascular adverse effects are a common cause of attrition in small molecule CNS programs, this improvement is highly significant in the context of long-term AD therapy.

In summary, among the three compounds evaluated, **G721-0377** exhibits the most favorable in vitro PK and safety profile for development as an AD therapeutic, with properties consistent with improved systemic stability, enhanced drug-like behavior, and a larger safety margin (Table 1). These results provide a strong rationale for prioritizing **G721-0377** for in vivo evaluation in appropriate AD models.

### Functional Validation of G721 Compounds as CHI3L1-Targeted AD Therapeutics

To evaluate whether the **G721** series can reverse CHI3L1-driven astrocytic dysfunction relevant to AD, we assessed their effects on amyloid uptake, lysosomal proteolytic activity, vesicular acidification, cytokine release, and inflammatory transcriptional signaling in human iPSC-derived astrocytes exposed to recombinant CHI3L1.

#### Reversal of CHI3L1-Induced Astrocytic Amyloid Dysfunction

As expected, treatment with CHI3L1 (300 ng/mL) significantly impaired uptake of pHrodo-Aβ_1-42_, reducing internalization to approximately two-thirds of vehicle control levels (Figure 5A). Neither **G721-0282** nor **G721-0179** significantly restored amyloid uptake at any concentration tested (10-50 µM). In contrast, **G721-0377** produced a clear, concentration-dependent rescue of endocytic function, achieving statistically significant recovery at 10 µM and near-complete normalization at 50 µM. As shown in Figure 5A, an anti-CHI3L1 antibody was used as a positive control. These data indicate that only **G721-0377** effectively reverses the CHI3L1-induced impairment in astrocytic amyloid handling. A parallel pattern was observed in lysosomal proteolysis. CHI3L1 markedly reduced DQ-BSA degradation, consistent with impaired lysosomal function (Figure 5B). Neither **G721-0282** nor **G721-0179** measurably corrected this defect. In contrast, **G721-0377** restored degradative capacity in a dose-dependent manner, returning lysosomal activity toward baseline at higher concentrations. Because defective lysosomal turnover is a central pathological feature of AD, the ability of **G721-0377** to normalize this pathway underscores its therapeutic relevance.

**Figure 5.**
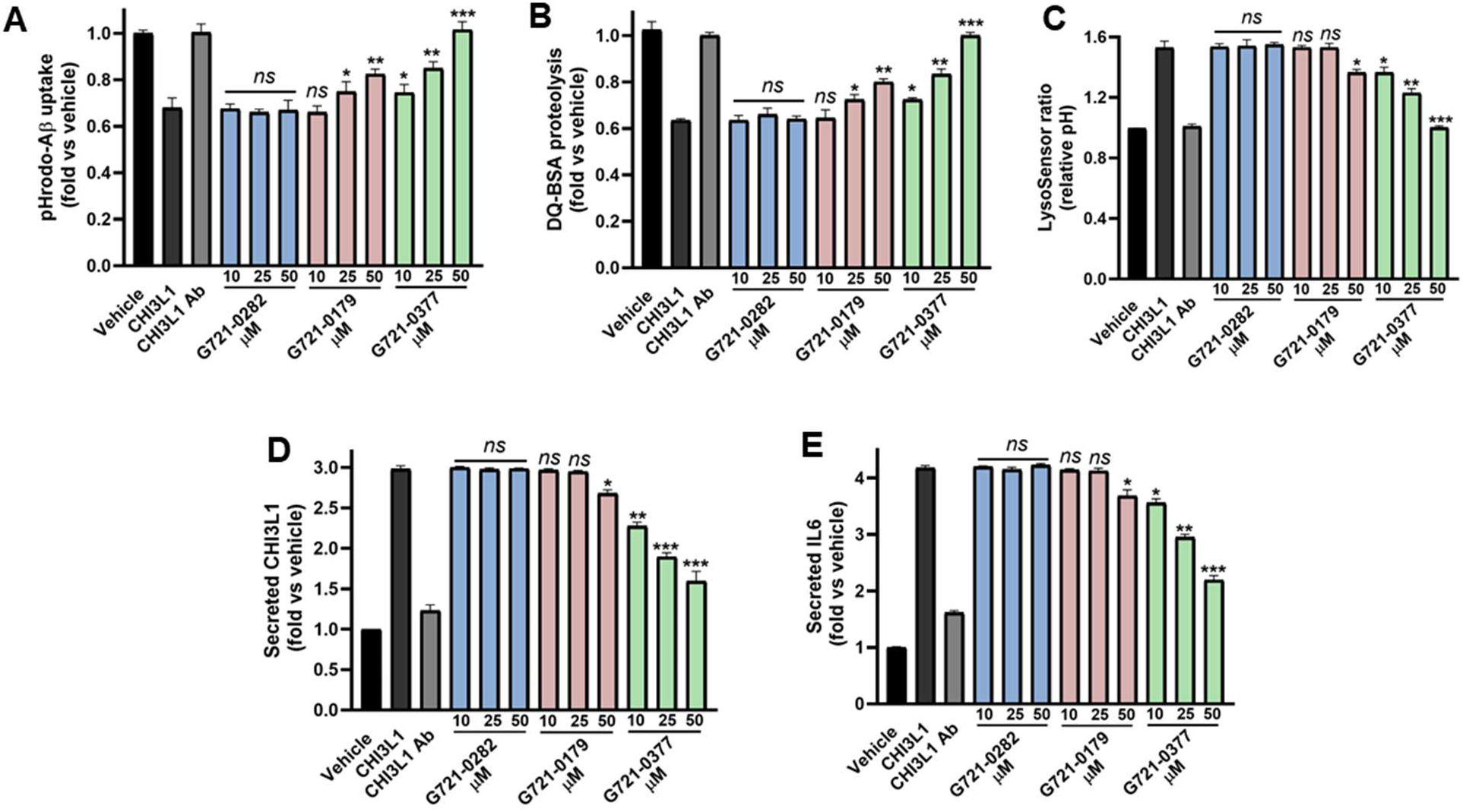
G721-0377 selectively restores amyloid uptake, lysosomal function, and cytokine balance in CHI3L1-treated astrocytes. Human iPSC-derived astrocytes were treated with recombinant CHI3L1 (300 ng/mL) and **G721** compounds (10-50 µM). Only **G721-0377** restored pHrodo-Aβ_1-42_ uptake (**A**), lysosomal proteolysis (DQ-BSA) (**B**), and vesicular acidification (LysoSensor) (**C**). G721-0377 also significantly suppressed CHI3L1 and IL-6 secretion (**D-E**), while **G721-0282** and **G721-0179** showed minimal activity. Anti-CHI3L1 antibody served as a positive control. One-way ANOVA with Dunnett’s post hoc test versus CHI3L1: ns, not significant; p < 0.05 (*), p < 0.01 (**), p < 0.001 (***).

#### Restoration of Lysosomal Acidification and Proteolytic Function

Disrupted lysosomal acidification further contributes to impaired proteostasis in neurodegeneration. LysoSensor analysis revealed that CHI3L1 significantly elevated lysosomal pH, indicating compromised vesicular acidification (Figure 5C). Neither **G721-0282** nor **G721-0179** reversed this effect. In contrast, **G721-0377** normalized lysosomal pH in a concentration-dependent manner, restoring acidification to near-vehicle levels at 50 µM. Together with the DQ-BSA data, these results demonstrate that **G721-0377** not only improves lysosomal activity but corrects the upstream physicochemical environment needed for proteolysis.

#### Selective Suppression of CHI3L1 and IL-6 Without Viability Loss

In addition to cellular clearance mechanisms, inflammatory output was evaluated. CHI3L1 dramatically increased secretion of both CHI3L1 and IL-6 (Figure 5D,E). Neither **G721-0282** nor **G721-0179** significantly reduced cytokine secretion across the concentration range studied. Conversely, **G721-0377** robustly suppressed release of both mediators in a dose-responsive manner, approaching baseline levels at higher concentrations. Importantly, none of the compounds affected astrocyte viability, confirming that the observed functional rescue was not attributable to cytotoxicity.

#### Suppression of CHI3L1-Induced NF-κB Activation

To determine whether these functional effects were accompanied by interruption of inflammatory signaling, we examined NF-κB transcriptional activity using a luciferase reporter assay. CHI3L1 induced a pronounced increase in NF-κB activity, consistent with its role as a pro-inflammatory effector (Figure 6A). **G721-0282** was inactive, and **G721-0179** produced only weak suppression at the highest dose tested. In contrast, **G721-0377** strongly and dose-dependently reduced NF-κB activation, reaching levels comparable to a neutralizing anti-CHI3L1 antibody. Viability controls confirmed that the observed suppression was not due to nonspecific effects (Figure 6B).

**Figure 6.**
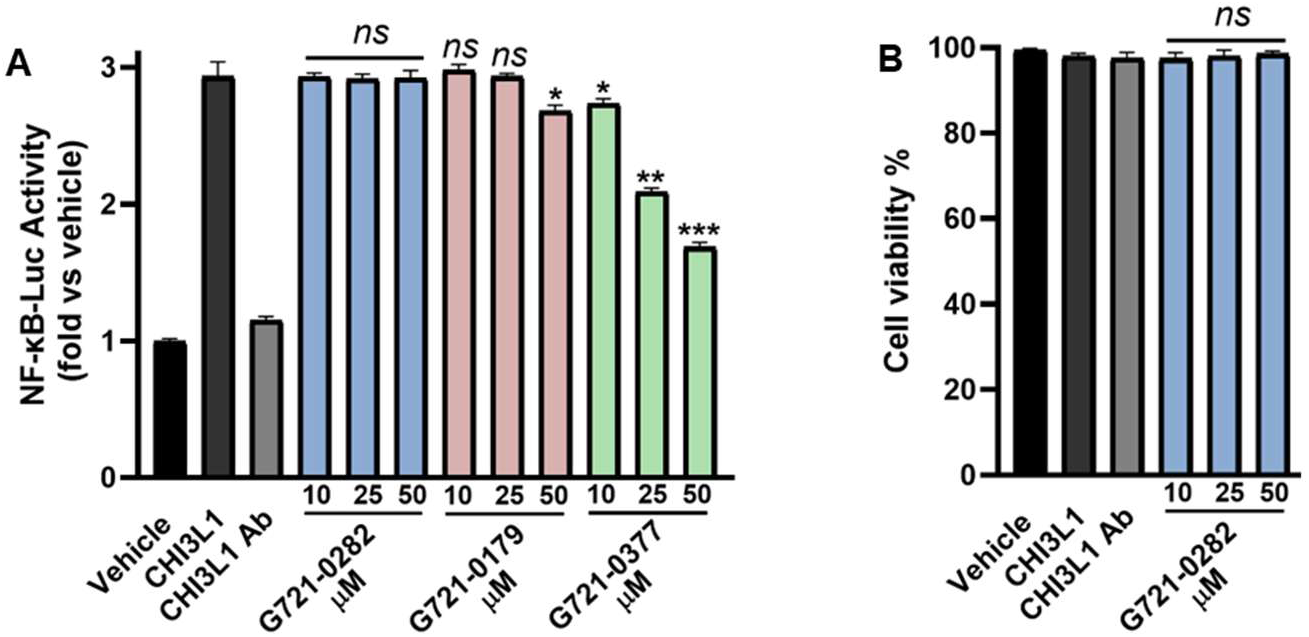
G721-0377 suppresses CHI3L1-induced NF-κB activation. **G721-0377** reduced NF-κB luciferase signaling in a dose-dependent manner, while **G721-0282** was inactive and **G721-0179** showed weak partial suppression only at 50 µM (A). No cytotoxicity was observed for **G721-0377** against the assay cells (B). Data represent mean ± SD from n = 3 experiments. One-way ANOVA with Dunnett’s post hoc test versus CHI3L1: ns, not significant; p < 0.05 (*), p < 0.01 (**), p < 0.001 (***).

Collectively, these results establish a clear functional hierarchy among the three analogs. Despite structural similarity and favorable physicochemical metrics, both **G721-0282** and **G721-0179** failed to reverse CHI3L1-induced astrocytic dysfunction or inflammatory signaling. In contrast, **G721-0377** consistently restored amyloid clearance, lysosomal integrity, and cytokine balance while suppressing the core inflammatory transcriptional program driven by CHI3L1. This coherence across multiple orthogonal readouts strongly supports effective pathway-level inhibition of CHI3L1 signaling by **G721-0377**. These findings have direct implications for AD therapy. Astrocytes are central regulators of both amyloid turnover and neuroinflammatory tone, and dysfunction in either process accelerates disease progression. By simultaneously restoring lysosomal competence and dampening inflammatory signaling, **G721-0377** addresses two interconnected disease mechanisms rather than targeting amyloid in isolation. This dual functional rescue profile more closely matches the biological complexity of AD and may provide superior therapeutic leverage over single-pathway approaches.

### Molecular Modeling

The molecular dynamics simulations and binding free energy analyses were carried out to assess the stability and interaction profiles of two CHI3L1 inhibitors, **G721-0377** and **G721-0282**. The structural analyses reveal that both compounds occupy the CHI3L1 active site through engagement with a conserved set of residues, including the catalytically important aromatic residues TRP99, TRP352, and TYR141, as well as hydrophobic residues LEU140, LEU356, and PHE261 (Figure 7 A-C). The RMSD trajectories (Panel D) demonstrated that all three systems, unbound CHI3L1, CHI3L1-**G721-0377** complex, and CHI3L1-**G721-0282** complex, were predicted to establish a stable equilibration during the 100 ns simulation period. The ligand-bound complexes exhibit RMSD values predominantly ranging between 1.0-1.5 Å, indicating robust structural stability throughout the simulation. Notably, only compound **G721-0377** was able to exhibit increased stability profile when compared to the unbound CHI3L1 protein which aligns with its observed experimental potency. Meanwhile, **G721-0282** was possessed a comparable RMSD profile to the unbound protein which suggests that ligand binding does not induce significant global conformational changes in CHI3L1 but rather stabilizes the existing binding pocket architecture. The MM-GBSA binding free energy calculations (Figure 7E) suggested the presence of a distinct energetic profiles for the two compounds. **G721-0377** (blue) demonstrates more favorable and consistent binding energies, with values frequently reaching -40 to -60 kcal/mol throughout the trajectory. In contrast, **G721-0282** (orange) exhibits greater fluctuation and generally less favorable binding energies, suggesting a more dynamic or less optimal binding mode. This energetic difference correlates with the observed experimental difference in the potency of the two compounds and further validate our findings. The protein-ligand contact frequency maps (Figure 7F-G) provide detailed characterization of the binding modes. **G721-0377** was predicted to establish sustained contacts with multiple residues throughout the simulation, as evidenced by the consistent contact patterns in the heatmap (Figure 7F). Key interactions are maintained with aromatic residues in the binding pocket, with notable engagement of residues in the 60-100 region of the protein sequence. The total contact plot shows sustained interaction density after an initial equilibration period around 40 ns, indicating stable ligand positioning. In comparison, **G721-0282** (Figure 7G) displays a weak contact profile with more pronounced variation in interaction patterns which corresponds with its observed lowered potency. Overall, these computational analyses suggest that while both compounds bind to the CHI3L1 active site and engage similar residues, **G721-0377** demonstrates superior binding characteristics, including more favorable binding energies, enhanced stability, and more consistent protein-ligand contacts throughout the simulation. The data indicate that **G721-0377** achieves a more optimal balance of electrostatic and hydrophobic interactions within the binding pocket, resulting in a thermodynamically and structurally more stable complex. These findings provide mechanistic insights that can inform structure-based optimization efforts for developing more potent CHI3L1 inhibitors and support our experimental findings.

**Figure 7.**
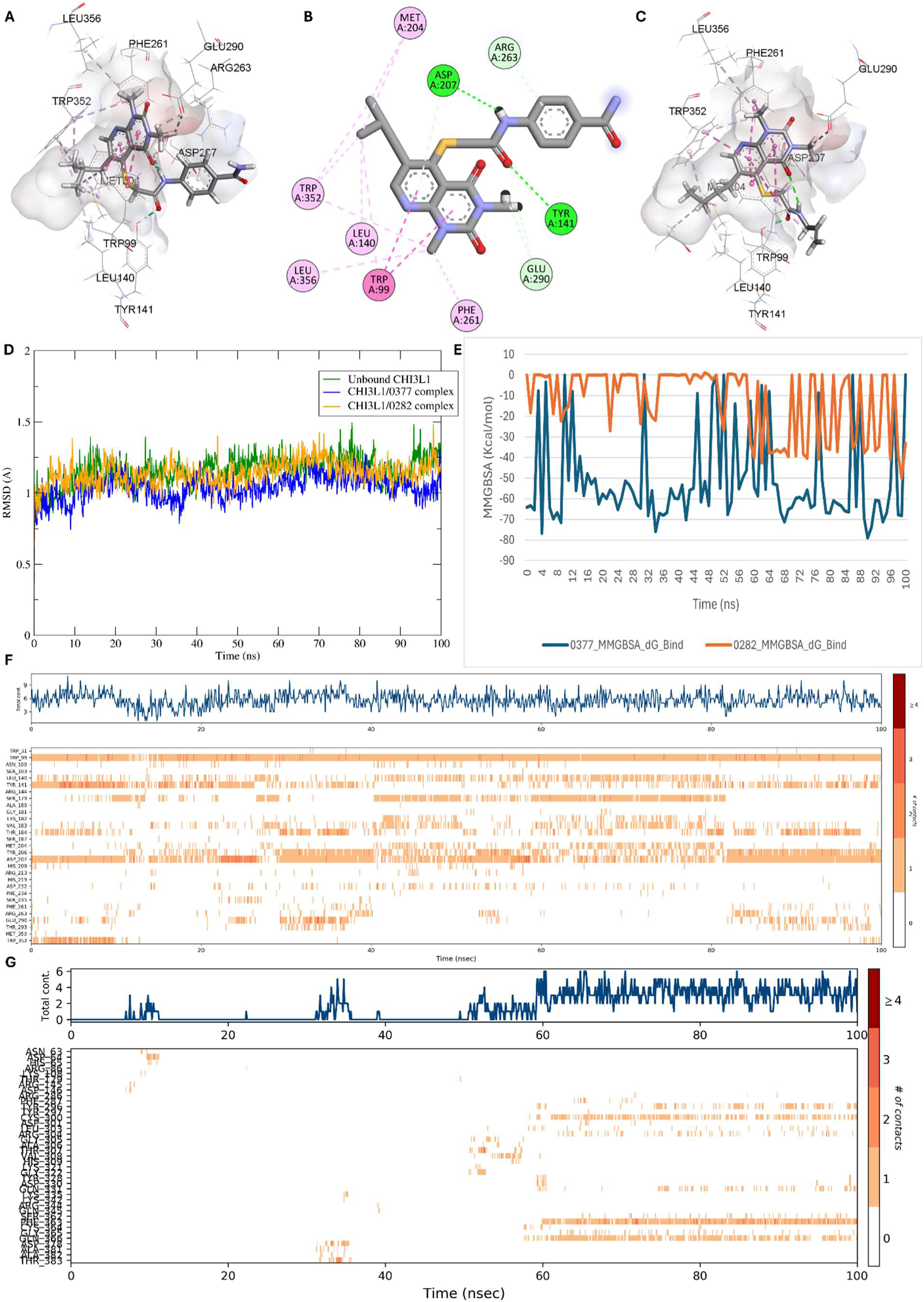
Molecular dynamics simulation and binding analysis of CHI3L1 inhibitors G721-0377 and G721-0282. (A-C) Three-dimensional representations of ligand-protein interactions. (A) Binding pose of **G721-0377** in the CHI3L1 active site. (B) Two-dimensional interaction diagram of **G721-0377**. (C) Binding pose of **G721-0282** in the CHI3L1 active site. (D) RMSD analysis over 100 ns molecular dynamics simulation for unbound CHI3L1 (green), **G721-0377** complex (blue), and **G721-0282**complex (orange). (E) MM-GBSA binding free energy calculations throughout the simulation trajectory for compounds **G721-0377** (blue) and **G721-0282** (orange), with time course shown in nanoseconds and energy in kcal/mol. (F) Protein-ligand contact analysis for **G721-0377** showing interaction frequency across the 100 ns simulation. Upper panel displays total contact count over time; lower heatmap illustrates per-residue contact frequencies with color intensity representing number of contacts. (G) Protein-ligand contact analysis for **G721-0282** with identical representation format, revealing distinct contact patterns and stability profile compared to **G721-0377**.

## Conclusions

This work demonstrates a rational strategy for optimizing small-molecule inhibition of CHI3L1, a key pathological mediator in Alzheimer’s disease. Virtual screening identified 24 analogues with promising predicted interactions, revealing clear structure-activity relationships that guided compound selection. Across all biophysical, pharmacokinetic, and functional evaluations, **G721-0377** emerged as the strongest candidate. It exhibited a significantly improved binding affinity (K_d_ = 45 µM), higher solubility (32.7 µM), balanced lipophilicity (logD₇.₄ = 3.8), reduced microsomal clearance (24–22 mL/min/mg), and a markedly safer hERG profile (IC50 = 18.2 µM). These improved properties translated directly into biological efficacy. **G721-0377** uniquely restored amyloid uptake, lysosomal proteolysis, and vesicular acidification, while robustly suppressing IL-6 and CHI3L1 secretion and inhibiting NF-κB-dependent inflammatory signaling. In contrast, **G721-0282** and **G721-0179** showed minimal activity despite structural similarity. By simultaneously correcting defects in both lysosomal homeostasis and inflammatory output, two processes central to AD progression, **G721-0377** addresses the biological complexity of AD beyond single-pathway targeting. The convergence of affinity, PK robustness, safety, and functional rescue strongly supports advancement of **G721-0377** into in vivo AD models and establishes a foundation for developing next-generation CHI3L1-targeted therapeutics.

## Experimental Section

### Evaluation of PK and physicochemical properties

The These experiments were conducted following our previously reported methods.^35^ The evaluations included LogD7.4 determination, microsomal stability, kinetic solubility, and cytotoxicity profiling across multiple cell lines. Solubility was assessed using UV–visible spectrophotometry, while cell viability was measured using the PrestoBlue assay.

### Cell culture and reagents

Human iPSC-derived astrocytes were maintained according to the manufacturer’s protocol in astrocyte medium (DMEM/F12 supplemented with 10% FBS) at 37 °C in a humidified atmosphere of 5% CO₂. Recombinant human CHI3L1 (YKL-40) was purchased from R&D Systems. For comparison and pathway validation, a neutralizing anti-CHI3L1 antibody (R&D Systems) was used as a positive control. All assays were performed in 96-well format unless otherwise indicated.

### Astrocyte Aβ Uptake and Lysosomal Function Assay

Human iPSC-astrocytes were seeded at 2 × 10^4^ cells per well (96-well plates) and cultured for 48 h. Cells were pre-treated with the tested compounds (10-50 µM, 30 min) prior to stimulation with CHI3L1 (300 ng mL⁻¹, 24 h). For Aβ uptake, cells were incubated with pHrodo™-Red–labeled Aβ_1-42_ (1 µM) for 2 h, dissociated using Accutase, fixed in 2% paraformaldehyde, and analyzed on a BD Fortessa flow cytometer (Ex/Em 560/585 nm). The mean fluorescence intensity (MFI) from ≥10,000 live events per well was normalized to vehicle controls.

Lysosomal degradation was assessed using DQ-BSA Green (10 µg mL⁻¹, 2 h), and lysosomal pH was measured using LysoSensor™ DND-160 (Ex/Em 340/440–535 nm ratio). Supernatants were collected for CHI3L1 and IL-6 quantification by ELISA (R&D Systems). Cell viability was assessed using CellTiter-Glo® (Promega). All measurements were performed in triplicate from at least three independent experiments.

### NF-κB Reporter Assay

To evaluate the effect of the **G721** compounds on CHI3L1-driven inflammatory signaling, human iPSC-astrocytes were transduced with an NF-κB-firefly luciferase reporter and a Renilla luciferase internal control. Cells were maintained for 72 h post-transduction to achieve stable expression before treatment. Cells were pre-incubated with the tested compounds (10-50 µM, 30 min) followed by stimulation with CHI3L1 (300 ng mL⁻¹, 6 h). Firefly and Renilla luciferase activities were measured using the Dual-Glo® Luciferase Assay System (Promega). Results were expressed as normalized Firefly/Renilla ratios and presented as fold-change relative to untreated vehicle controls. Cell viability was verified in parallel using CellTiter-Glo®.

### MST Screening

Protein–ligand interactions were assessed by microscale thermophoresis (MST) following our previously established protocol. CHI3L1-His was fluorescently labeled with RED-tris-NTA dye (NanoTemper Technologies) according to the manufacturer’s guidelines. The labeled protein was then incubated with test compounds under optimized assay conditions, and measurements were performed using the Dianthus NT.23 Pico system.

To minimize assay artifacts caused by compound-related autofluorescence or fluorescence quenching, dedicated control experiments were included. For autofluorescence assessment, the fluorescence of labeled protein in 2.5% DMSO was compared with buffer containing 250 μM compound in 2.5% DMSO (Figure S1A). To evaluate quenching, fluorescence from 20 nM dye in buffer with 2.5% DMSO was compared with the fluorescence of 20 nM dye incubated with each compound under identical solvent conditions (Figure S1B). Compounds showing fluorescence values outside the mean ± 3 SD of the corresponding controls were excluded from further analysis.

### Molecular Modeling

The three-dimensional structures of CHI3L1-inhibitor complexes were prepared using Schrödinger suite (Schrödinger, LLC, New York, NY) with protein preparation and ligand docking performed to generate initial binding poses for G721-0377 and G721-0282 at the default settings. Molecular dynamics simulations were conducted using DESMOND molecular dynamics package over a 100 ns trajectory using the same protocol employed in our previous work.^36,43^ All structural visualizations, including three-dimensional binding poses and two-dimensional interaction diagrams, were generated using Discovery Studio Visualizer (BIOVIA, Dassault Systèmes), which was employed to illustrate hydrogen bonding patterns, hydrophobic contacts, and other non-covalent interactions between the inhibitors and CHI3L1 binding pocket residues.

## Acknowledgments

This work was supported by the National Institute of Neurological Disorders and Stroke under grant number R01NS136524 (PI: Gabr).

## Abbreviations

CHI3L1: Chitinase-3-like protein 1
MST: Microscale thermophoresis
SPR: Surface plasmon resonance
Kd: Equilibrium dissociation constant
Aβ: Amyloid-β
CSF: Cerebrospinal fluid
MAPK: Mitogen-activated protein kinase
STAT3: Signal transducer and activator of transcription 3
PK: Pharmacokinetics
BBB: Blood–brain barrier
NF-κB: Nuclear factor kappa-light-chain-enhancer of activated B cells
PAMPA: Parallel artificial membrane permeability assay
CNS: Central nervous system
DQ-BSA: Dye-quenched bovine serum albumin
IL-6: Interleukin-6
Human iPSC: Human induced pluripotent stem cells

---

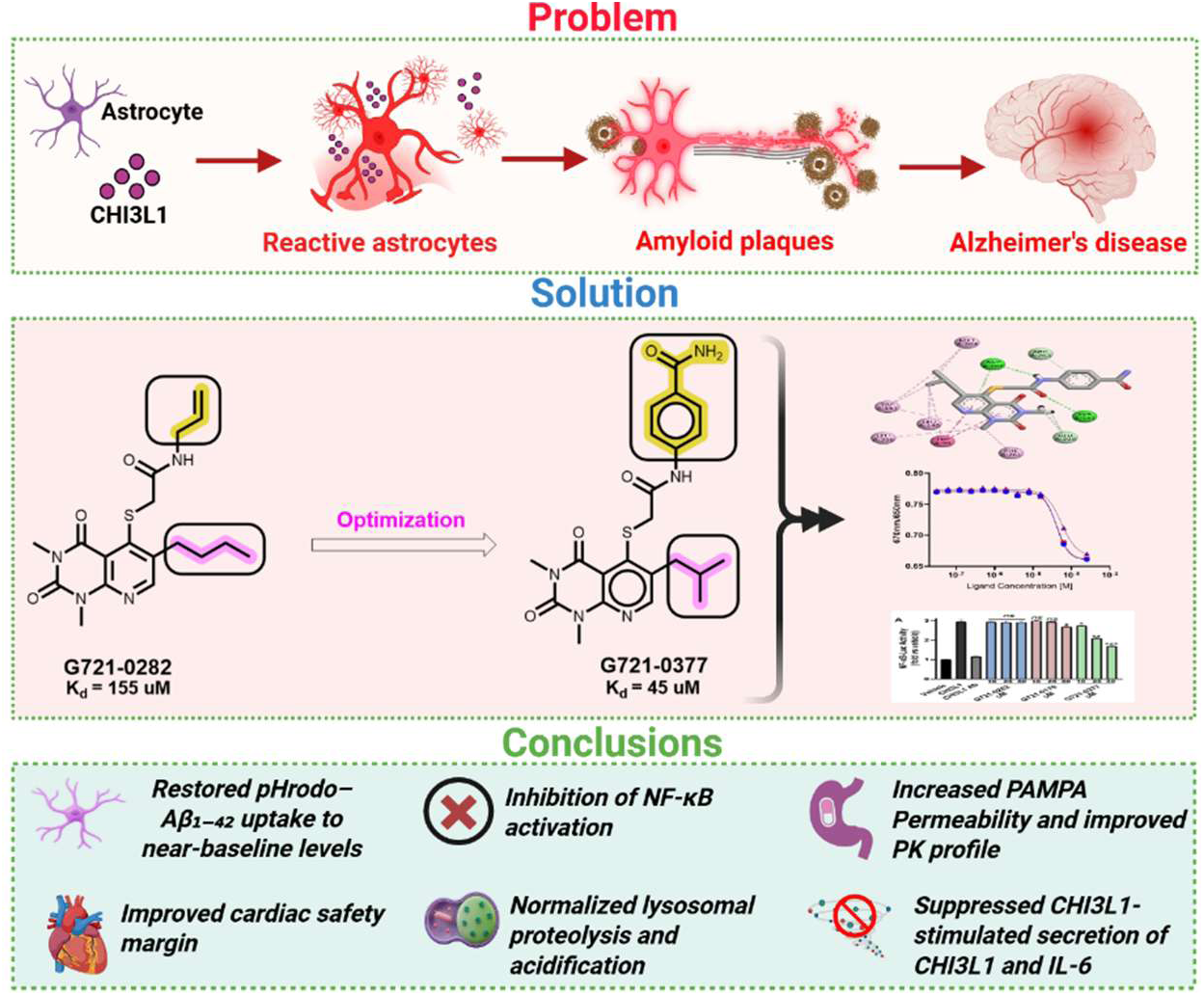

