## Supporting Information for "Targeting CHI3L1 in Alzheimer’s Disease: Optimization of G721-0282 and Functional Evaluation in Astrocyte Models"

* To whom correspondence should be addressed:

**Table of Contents**

| 1. Smiles of 24 hit compounds | S1 |
| --- | --- |
| 3. MST studies 2. Control experiments | S2 |
| 3. Docking scores | S3 |

**1. Smiles of 24 hit compounds**

| **S. No.** | **IDNUMBER** | **SMILES** |
| --- | --- | --- |
| 1 | G721-0134 | CC(C)c(c(SCC(NCC=C)=O)c1C(N2C)=O)cnc1N(C)C2=O |
| 2 | G721-0176 | CCCCc(c(SCC(Nc1cccc2ccccc12)=O)c1C(N2C)=O)cnc1N(C)C2=O |
| 3 | G721-0179 | CCCCc(c(SCC(Nc1ccc(C)cc1)=O)c1C(N2C)=O)cnc1N(C)C2=O |
| 4 | G721-0211 | CCCCc(c(SCC(Nc(cc1)ccc1NC(C)=O)=O)c1C(N2C)=O)cnc1N(C)C2=O |
| 5 | G721-0217 | CCCCc(c(SCC(NCc(cc1)cc2c1OCO2)=O)c1C(N2C)=O)cnc1N(C)C2=O |
| 6 | G721-0226 | CCCCc(c(SCC(Nc1nnc(SCC)s1)=O)c1C(N2C)=O)cnc1N(C)C2=O |
| 7 | G721-0275 | CCCCc(c(SCC(NCCC)=O)c1C(N2C)=O)cnc1N(C)C2=O |
| 8 | G721-0278 | CCCCc(c(SCC(NCCOC)=O)c1C(N2C)=O)cnc1N(C)C2=O |
| 9 | G721-0325 | CC(C)Cc(c(SCC(Nc1c(C)cccc1)=O)c1C(N2C)=O)cnc1N(C)C2=O |
| 10 | G721-0336 | CC(C)Cc(c(SCC(Nc(cc1)ccc1F)=O)c1C(N2C)=O)cnc1N(C)C2=O |
| 11 | G721-0346 | CC(C)Cc(c(SCC(Nc(cc1)cc2c1OCO2)=O)c1C(N2C)=O)cnc1N(C)C2=O |
| 12 | G721-0362 | CCOC(CNC(CSc(c(CC(C)C)cnc1N(C)C(N2C)=O)c1C2=O)=O)=O |
| 13 | G721-0365 | CC(C)Cc(c(SCC(NCc(cc1)cc2c1OCO2)=O)c1C(N2C)=O)cnc1N(C)C2=O |
| 14 | G721-0366 | CC(C)Cc(c(SCC(NCc1ccco1)=O)c1C(N2C)=O)cnc1N(C)C2=O |
| 15 | G721-0377 | CC(C)Cc(c(SCC(Nc(cc1)ccc1C(N)=O)=O)c1C(N2C)=O)cnc1N(C)C2=O |
| 16 | G721-0430 | CC(C)Cc(c(SCC(NCC=C)=O)c1C(N2C)=O)cnc1N(C)C2=O |
| 17 | G721-0510 | CCOC(CNC(CSc(c(COC)cnc1N(C)C(N2C)=O)c1C2=O)=O)=O |
| 18 | G721-0513 | CN(c1ncc(COC)c(SCC(NCc(cc2)cc3c2OCO3)=O)c1C(N1C)=O)C1=O |
| 19 | G721-0525 | Cc1ccc(CNC(CSc(c(COC)cnc2N(C)C(N3C)=O)c2C3=O)=O)cc1 |
| 20 | G721-0579 | CN(c1ncc(COC)c(SCC(NCC=C)=O)c1C(N1C)=O)C1=O |
| 21 | G857-1029 | CCCc(c(SCC(Nc1cccc2ccccc12)=O)c1C(N2C)=O)cnc1N(C)C2=O |
| 22 | G857-1036 | CCCc(c(SCC(Nc1cc(C)cc(C)c1)=O)c1C(N2C)=O)cnc1N(C)C2=O |
| 23 | G857-1053 | CCCc(c(SCC(Nc(cc1)ccc1C(N)=O)=O)c1C(N2C)=O)cnc1N(C)C2=O |
| 24 | G857-1069 | CCCc(c(SCC(NCC=C)=O)c1C(N2C)=O)cnc1N(C)C2=O |

**2. Control experiments**

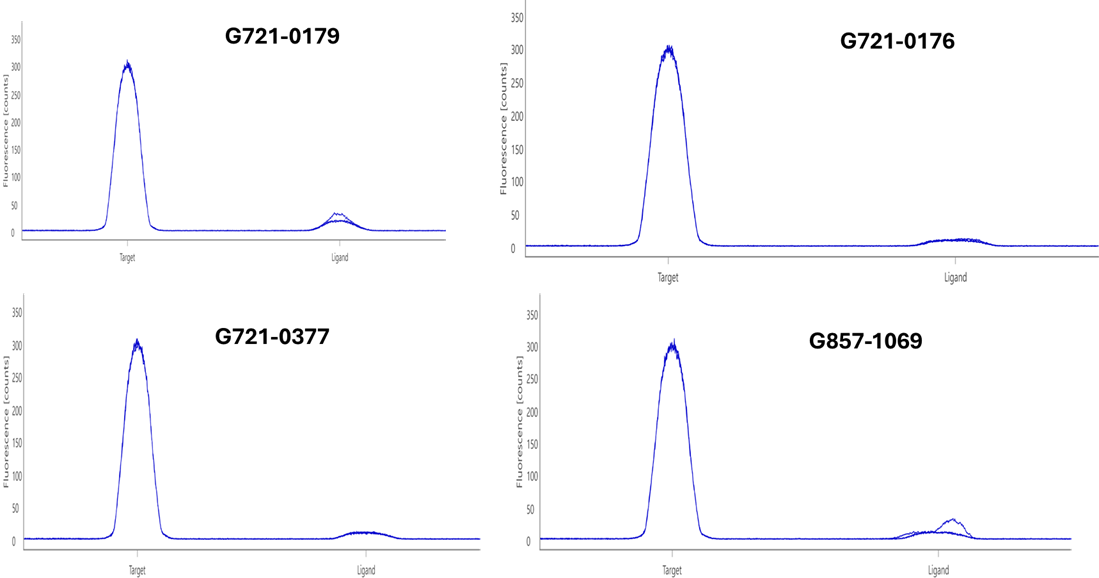

**Figure S1.** Assessment of potential artifacts from autofluorescence by comparing the fluorescence of labeled protein in 2.5% DMSO with buffer containing 250 μM compound in 2.5% DMSO.

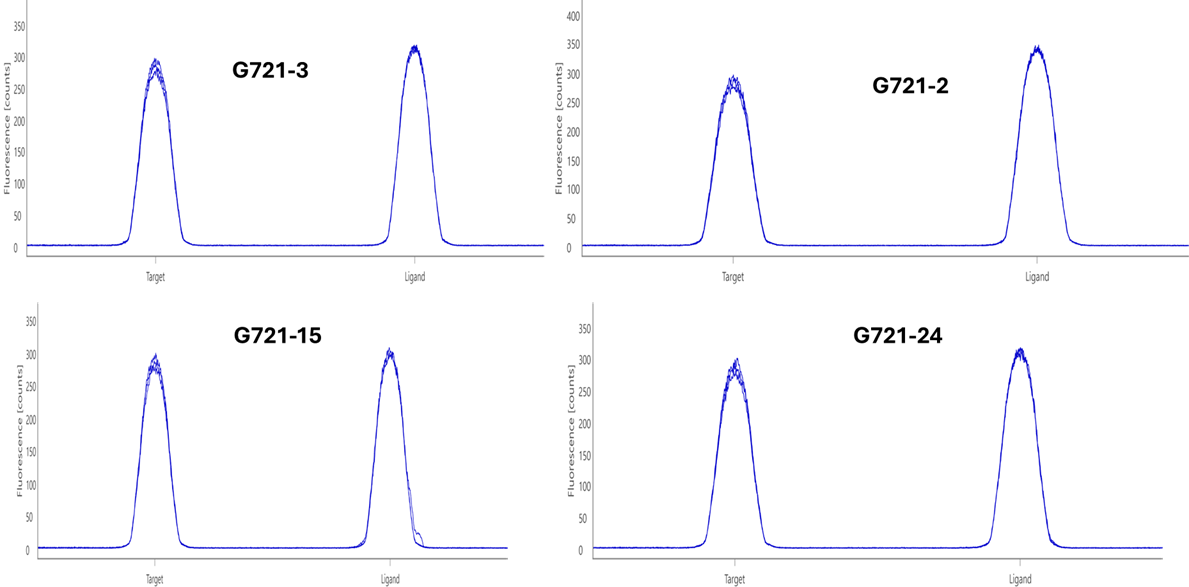

**Figure S2**. Evaluation of potential quenching-related artifacts in MST assays. Fluorescence of 20 nM dye in 2.5% DMSO was compared with that of 20 nM dye incubated with 250 μM compound in 2.5% DMSO to assess compound-induced quenching effects.

**3. Docking Scores**

**Table S1. Docking Scores of commercially available G721-0282 analogues**

| **Id number** | **Docking_score** | **Id number** | **Docking_score** | **Id number** | **Docking_score** |
| --- | --- | --- | --- | --- | --- |
| G721-0001F | -6.55313 | G721-0233 | -3.40346 | G721-0533 | -5.62166 |
| G721-0005 | -6.15165 | G721-0234 | -5.39958 | G721-0534 | -7.15551 |
| G721-0007 | -4.84031 | G721-0235 | -6.46598 | G721-0536 | -6.06612 |
| G721-0009 | -7.06265 | G721-0236 | -6.46757 | G721-0537 | -5.03923 |
| G721-0015 | -5.9598 | G721-0237 | -5.90653 | G721-0538 | -5.42724 |
| G721-0016 | -5.92489 | G721-0239 | -4.30831 | G721-0539 | -5.23266 |
| G721-0017 | -5.54368 | G721-0240 | -4.33954 | G721-0540 | -6.32064 |
| G721-0018 | -7.00597 | G721-0241 | -6.78145 | G721-0542 | -5.7104 |
| G721-0019 | -5.39256 | G721-0242 | -4.68419 | G721-0543 | -4.96942 |
| G721-0020 | -5.89212 | G721-0243 | -5.0335 | G721-0545 | -6.23846 |
| G721-0022 | -5.47988 | G721-0245 | -4.44225 | G721-0547 | -6.92629 |
| G721-0023 | -4.22188 | G721-0246 | -4.87661 | G721-0548 | -5.65606 |
| G721-0024 | -4.04296 | G721-0247 | -8.123 | G721-0549 | -6.21349 |
| G721-0025 | -5.66328 | G721-0248 | -6.23828 | G721-0550 | -6.16863 |
| G721-0026 | -4.72549 | G721-0249 | -6.41529 | G721-0560 | -6.73332 |
| G721-0027 | -5.79598 | G721-0250 | -7.39142 | G721-0565 | -4.83001 |
| G721-0028 | -4.51911 | G721-0251 | -6.34936 | G721-0569 | -6.13222 |
| G721-0029 | -4.61921 | G721-0252 | -7.02242 | G721-0571F | -6.20097 |
| G721-0030 | -5.33093 | G721-0263 | -7.57046 | G721-0572 | -5.05214 |
| G721-0031 | -5.82956 | G721-0268 | -5.56395 | G721-0576 | -5.35136 |
| G721-0032 | -5.25782 | G721-0272 | -5.34213 | G721-0578 | -6.20202 |
| G721-0033 | -6.97991 | G721-0274 | -8.20397 | G721-0579 | -6.32618 |
| G721-0034 | -4.12161 | G721-0275 | -8.57286 | G721-0580 | -5.42411 |
| G721-0035 | -4.59578 | G721-0276 | -8.16697 | G721-0581 | -4.63162 |
| G721-0036 | -4.62319 | G721-0277 | -4.49609 | G721-0583 | -5.97024 |
| G721-0037 | -5.29977 | G721-0278 | -9.29903 | G721-0585 | -6.21725 |
| G721-0038 | -5.0586 | G721-0280 | -6.08163 | G721-0586 | -7.7019 |
| G721-0039 | -5.27136 | G721-0280 | -5.52194 | G721-0587 | -5.97479 |
| G721-0040 | -5.36381 | G721-0280 | -4.29461 | G721-0588 | -6.96339 |
| G721-0041 | -5.33638 | G721-0281 | -5.8798 | G721-0589 | -5.95583 |
| G721-0042 | -6.60366 | G721-0283 | -3.61036 | G721-0590 | -7.06355 |
| G721-0043 | -6.72838 | G721-0284 | -4.58641 | G721-0591 | -7.34122 |
| G721-0044 | -4.85952 | G721-0286 | -3.44725 | G721-0593 | -5.74248 |
| G721-0045 | -5.74375 | G721-0287 | -7.83865 | G786-1490 | -7.38572 |
| G721-0046 | -5.45839 | G721-0288 | -7.86146 | G857-0877 | -7.62355 |
| G721-0048 | -5.62446 | G721-0289 | -6.1622 | G857-0880 | -6.40628 |
| G721-0049 | -5.35009 | G721-0291 | -6.17316 | G857-0881 | -6.39814 |
| G721-0051 | -5.44826 | G721-0292 | -4.68151 | G857-0882 | -5.28883 |
| G721-0052 | -6.52179 | G721-0293 | -6.66582 | G857-0883 | -5.35403 |
| G721-0053 | -6.13036 | G721-0294 | -5.56442 | G857-0884 | -5.41841 |
| G721-0054 | -6.77921 | G721-0295 | -5.47555 | G857-0885 | -5.43564 |
| G721-0055 | -4.54796 | G721-0296 | -6.52463 | G857-0886 | -5.58111 |
| G721-0056 | -3.96427 | G721-0306 | -7.05245 | G857-0887 | -5.58924 |
| G721-0057 | -5.46913 | G721-0312 | -4.36402 | G857-0888 | -6.41496 |
| G721-0058 | -5.40038 | G721-0313 | -5.92787 | G857-0889 | -5.01088 |
| G721-0059 | -6.04047 | G721-0314 | -4.4581 | G857-0890 | -4.71332 |
| G721-0060 | -6.56991 | G721-0315 | -4.43133 | G857-0891 | -6.05255 |
| G721-0062 | -7.87158 | G721-0316 | -5.67011 | G857-0892 | -5.81055 |
| G721-0063 | -6.36683 | G721-0321 | -4.41983 | G857-0893 | -5.84867 |
| G721-0064 | -4.07704 | G721-0322 | -6.01699 | G857-0894 | -5.05835 |
| G721-0065 | -4.75436 | G721-0323 | -7.5168 | G857-0895 | -5.62121 |
| G721-0066 | -7.60768 | G721-0324 | -4.17419 | G857-0896 | -3.95929 |
| G721-0067 | -6.90725 | G721-0325 | -8.64253 | G857-0897 | -6.89492 |
| G721-0068 | -6.66509 | G721-0326 | -8.01358 | G857-0898 | -5.70705 |
| G721-0069 | -8.19739 | G721-0327 | -3.72565 | G857-0899 | -6.28485 |
| G721-0071 | -7.84927 | G721-0328 | -5.27719 | G857-0900 | -7.13722 |
| G721-0072 | -8.53527 | G721-0329 | -5.62028 | G857-0901 | -6.18955 |
| G721-0073 | -4.86895 | G721-0330 | -5.81193 | G857-0903 | -5.63042 |
| G721-0073 | -3.39466 | G721-0331 | -5.5209 | G857-0904 | -7.03857 |
| G721-0074 | -5.68171 | G721-0332 | -6.19184 | G857-0905 | -6.85072 |
| G721-0074 | -2.01136 | G721-0333 | -8.14995 | G857-0906 | -5.22082 |
| G721-0075 | -5.96678 | G721-0334 | -6.21128 | G857-0909 | -6.54816 |
| G721-0076 | -4.35982 | G721-0336 | -8.81405 | G857-0910F | -5.60674 |
| G721-0077 | -4.83966 | G721-0337 | -8.3555 | G857-0911 | -6.03162 |
| G721-0077 | -3.91264 | G721-0338 | -8.42802 | G857-0912 | -4.99201 |
| G721-0078 | -3.68332 | G721-0339 | -8.5339 | G857-0913 | -6.02426 |
| G721-0078 | -3.5193 | G721-0340 | -5.89083 | G857-0914 | -6.63563 |
| G721-0079 | -5.41778 | G721-0341 | -5.32774 | G857-0915 | -6.02266 |
| G721-0080 | -5.12814 | G721-0342 | -4.94831 | G857-0916 | -6.57132 |
| G721-0081 | -6.99535 | G721-0343 | -8.79875 | G857-0919 | -6.06566 |
| G721-0082 | -4.89361 | G721-0344 | -4.54302 | G857-0920 | -6.22858 |
| G721-0084 | -3.40588 | G721-0345 | -7.99885 | G857-0921 | -5.22882 |
| G721-0084 | -4.7961 | G721-0346 | -8.85061 | G857-0922 | -4.63455 |
| G721-0085 | -5.76382 | G721-0348 | -5.90642 | G857-0923 | -6.70474 |
| G721-0085 | -2.87279 | G721-0349 | -3.92194 | G857-0924 | -6.26211 |
| G721-0086 | -6.05014 | G721-0350 | -6.0892 | G857-0925 | -6.45597 |
| G721-0087 | -4.71666 | G721-0351 | -6.73957 | G857-0926 | -5.27171 |
| G721-0088 | -6.38495 | G721-0352 | -7.90334 | G857-0927 | -6.77787 |
| G721-0089 | -6.43694 | G721-0353 | -6.51264 | G857-0928 | -4.91236 |
| G721-0091 | -5.53911 | G721-0354 | -5.42647 | G857-0928 | -5.14435 |
| G721-0092 | -4.96072 | G721-0355 | -7.43423 | G857-0929 | -6.81729 |
| G721-0093 | -4.61959 | G721-0357 | -6.76231 | G857-0930 | -6.06388 |
| G721-0094 | -5.38633 | G721-0359 | -8.41776 | G857-0931 | -6.43817 |
| G721-0095 | -5.16405 | G721-0360 | -7.66925 | G857-0932 | -6.01928 |
| G721-0097 | -5.8768 | G721-0362 | -9.04236 | G857-0933 | -6.09527 |
| G721-0098 | -4.98082 | G721-0363 | -8.58729 | G857-0934 | -6.87524 |
| G721-0099 | -5.1584 | G721-0364 | -8.54246 | G857-0935 | -6.1828 |
| G721-0100 | -4.38592 | G721-0365 | -9.09781 | G857-0936 | -5.01767 |
| G721-0101 | -5.66157 | G721-0366 | -8.04234 | G857-0937 | -6.16635 |
| G721-0102 | -7.14744 | G721-0367 | -5.86163 | G857-0938 | -6.70358 |
| G721-0103 | -5.97009 | G721-0367 | -7.39436 | G857-0939 | -5.94738 |
| G721-0104 | -5.45413 | G721-0368 | -6.55149 | G857-0940 | -6.30852 |
| G721-0105 | -4.93484 | G721-0370 | -4.0272 | G857-0941 | -6.13351 |
| G721-0106 | -6.62724 | G721-0370 | -2.66754 | G857-0942 | -7.27104 |
| G721-0112 | -6.99043 | G721-0371 | -4.26003 | G857-0943 | -6.3459 |
| G721-0116 | -5.73434 | G721-0372 | -8.3446 | G857-0944 | -5.66133 |
| G721-0120 | -5.41712 | G721-0373 | -7.73639 | G857-0945 | -6.28112 |
| G721-0124 | -5.7576 | G721-0373 | -7.49856 | G857-0946 | -5.78942 |
| G721-0127 | -4.89635 | G721-0374 | -4.8306 | G857-0947 | -6.27821 |
| G721-0128 | -5.53053 | G721-0374 | -6.34448 | G857-0948 | -6.40205 |
| G721-0129 | -4.72959 | G721-0375 | -6.47167 | G857-0949 | -7.68087 |
| G721-0130 | -7.10191 | G721-0377 | -8.51635 | G857-0950 | -6.44931 |
| G721-0132 | -5.29446 | G721-0380 | -6.26463 | G857-0951 | -6.76004 |
| G721-0132 | -5.61807 | G721-0380 | -2.66476 | G857-0952 | -6.09908 |
| G721-0132 | -5.56001 | G721-0381 | -5.29892 | G857-0953 | -5.30571 |
| G721-0133 | -5.50738 | G721-0381 | -3.83764 | G857-0954 | -6.40253 |
| G721-0134 | -5.0055 | G721-0382 | -6.50704 | G857-0955 | -7.5227 |
| G721-0135 | -5.18414 | G721-0383 | -5.53842 | G857-0956 | -7.61752 |
| G721-0136 | -5.71422 | G721-0384 | -6.54423 | G857-0957 | -4.65228 |
| G721-0138 | -6.30611 | G721-0385 | -6.3001 | G857-0958 | -5.81332 |
| G721-0139 | -4.94299 | G721-0387 | -5.27068 | G857-0959 | -5.80276 |
| G721-0140 | -6.3483 | G721-0388 | -4.8508 | G857-0960 | -7.20496 |
| G721-0141 | -6.95885 | G721-0389 | -5.88152 | G857-0961 | -5.56834 |
| G721-0143 | -5.96982 | G721-0390 | -5.03917 | G857-0962 | -6.25859 |
| G721-0144 | -5.88939 | G721-0391 | -4.96886 | G857-0963 | -6.35486 |
| G721-0145 | -5.92347 | G721-0393 | -5.59359 | G857-0964 | -7.80111 |
| G721-0146 | -5.24419 | G721-0395 | -5.29662 | G857-0965 | -6.79728 |
| G721-0147 | -5.41974 | G721-0397 | -5.4753 | G857-0968 | -7.38715 |
| G721-0148 | -5.60649 | G721-0398 | -5.51622 | G857-0968 | -4.33481 |
| G721-0149 | -5.94921 | G721-0399 | -5.29572 | G857-0969 | -7.00007 |
| G721-0150 | -7.40758 | G721-0400 | -3.96894 | G857-0970 | -6.22728 |
| G721-0151 | -7.3343 | G721-0402 | -7.40772 | G857-0970 | -2.38901 |
| G721-0152 | -7.68049 | G721-0408 | -7.43304 | G857-0971 | -6.0927 |
| G721-0153 | -5.31081 | G721-0416 | -6.37803 | G857-0972 | -4.72647 |
| G721-0154 | -8.107 | G721-0420 | -5.32754 | G857-0973 | -6.3485 |
| G721-0156 | -8.21834 | G721-0421 | -6.83754 | G857-0974 | -5.03734 |
| G721-0158 | -7.3287 | G721-0422 | -4.81442 | G857-0975 | -6.55063 |
| G721-0159 | -7.54387 | G721-0423 | -7.25698 | G857-0976 | -6.60709 |
| G721-0165 | -4.21617 | G721-0424 | -8.19361 | G857-0977 | -5.63649 |
| G721-0166 | -6.12027 | G721-0425 | -4.76751 | G857-0978 | -5.18813 |
| G721-0169 | -5.25104 | G721-0426 | -6.27318 | G857-0979 | -5.97711 |
| G721-0171 | -6.59138 | G721-0427 | -3.93381 | G857-0980 | -5.3388 |
| G721-0172 | -3.42802 | G721-0428 | -3.99002 | G857-0981 | -5.88673 |
| G721-0173 | -4.49786 | G721-0428 | -5.6167 | G857-0982 | -6.0136 |
| G721-0174 | -8.03194 | G721-0428 | -6.25684 | G857-0983 | -4.33068 |
| G721-0175 | -3.76514 | G721-0429 | -5.8168 | G857-0984 | -5.78606 |
| G721-0176 | -8.40201 | G721-0430 | -8.51734 | G857-0985 | -5.41479 |
| G721-0177 | -6.28183 | G721-0431 | -5.46646 | G857-0986 | -7.00897 |
| G721-0178 | -4.29268 | G721-0432 | -5.43785 | G857-0987 | -7.81384 |
| G721-0179 | -8.61652 | G721-0434 | -7.99431 | G857-0988 | -6.36422 |
| G721-0180 | -5.65823 | G721-0435 | -8.4442 | G857-0989 | -6.51289 |
| G721-0181 | -4.11342 | G721-0436 | -8.42585 | G857-0990 | -5.71913 |
| G721-0182 | -6.42566 | G721-0437 | -5.30656 | G857-0991 | -4.8985 |
| G721-0183 | -5.8369 | G721-0438 | -8.71956 | G857-0992 | -5.5127 |
| G721-0184 | -6.39951 | G721-0439 | -4.21546 | G857-0992 | -5.76781 |
| G721-0185 | -8.77623 | G721-0445 | -4.19328 | G857-0993 | -5.34239 |
| G721-0186 | -7.9807 | G721-0462 | -4.74053 | G857-0993 | -5.97841 |
| G721-0187 | -5.7241 | G721-0469 | -5.82771 | G857-0994 | -5.67403 |
| G721-0188 | -5.32944 | G721-0470 | -6.38177 | G857-0994 | -5.83098 |
| G721-0189 | -4.90121 | G721-0471 | -6.03331 | G857-0995 | -7.53953 |
| G721-0190 | -4.99633 | G721-0472 | -7.82169 | G857-0996 | -5.5214 |
| G721-0191 | -5.62808 | G721-0473 | -5.6936 | G857-0997 | -4.81691 |
| G721-0192 | -8.33711 | G721-0474 | -5.63435 | G857-0997 | -4.92293 |
| G721-0193 | -5.48252 | G721-0475 | -6.38975 | G857-0997 | -4.33257 |
| G721-0194 | -7.81497 | G721-0476 | -5.43708 | G857-0998 | -5.91096 |
| G721-0195 | -7.78275 | G721-0478 | -5.1366 | G857-0999 | -6.92514 |
| G721-0196 | -4.94134 | G721-0479 | -5.6711 | G857-1000 | -6.18004 |
| G721-0197 | -8.70941 | G721-0480 | -6.02305 | G857-1001 | -5.01034 |
| G721-0198 | -7.98105 | G721-0481 | -6.58192 | G857-1002 | -5.82467 |
| G721-0199 | -6.77022 | G721-0482 | -5.98652 | G857-1004 | -4.60781 |
| G721-0200 | -6.25998 | G721-0484 | -6.84275 | G857-1005 | -6.21672 |
| G721-0201 | -3.82399 | G721-0485 | -4.77309 | G857-1006 | -7.51165 |
| G721-0202 | -6.7227 | G721-0486 | -5.54882 | G857-1007 | -5.9197 |
| G721-0203 | -7.98744 | G721-0488 | -4.58433 | G857-1008 | -6.37447 |
| G721-0204 | -3.99462 | G721-0489 | -6.36979 | G857-1009 | -5.8893 |
| G721-0205 | -7.52103 | G721-0490 | -5.54085 | G857-1010 | -5.90715 |
| G721-0206 | -4.84909 | G721-0491 | -5.66587 | G857-1011 | -5.86923 |
| G721-0207 | -8.00864 | G721-0492 | -4.97364 | G857-1013 | -5.56549 |
| G721-0208 | -9.02323 | G721-0493 | -4.8917 | G857-1014 | -5.11176 |
| G721-0209 | -7.46836 | G721-0496 | -7.8419 | G857-1015 | -5.8851 |
| G721-0210 | -5.73258 | G721-0497 | -6.77812 | G857-1016 | -5.523 |
| G721-0211 | -8.6947 | G721-0498 | -4.81542 | G857-1017 | -5.58589 |
| G721-0212 | -7.46158 | G721-0499 | -6.56222 | G857-1018 | -6.73906 |
| G721-0213 | -7.97441 | G721-0502 | -5.20857 | G857-1019 | -5.9157 |
| G721-0214 | -6.10264 | G721-0503 | -6.01899 | G857-1021 | -6.34793 |
| G721-0215 | -6.40403 | G721-0504 | -7.37794 | G857-1022 | -4.97734 |
| G721-0216 | -5.44207 | G721-0505 | -7.25165 | G857-1022 | -7.26552 |
| G721-0217 | -9.29193 | G721-0506 | -4.70527 | G857-1023 | -6.85124 |
| G721-0218 | -8.18896 | G721-0507 | -5.42157 | G857-1024 | -7.31246 |
| G721-0219 | -5.81067 | G721-0508 | -7.27395 | G857-1027 | -7.26154 |
| G721-0219 | -8.24999 | G721-0509 | -5.66653 | G857-1028 | -5.19254 |
| G721-0220 | -8.48319 | G721-0510 | -9.53906 | G857-1029 | -4.54083 |
| G721-0221 | -8.36859 | G721-0512 | -5.95339 | G857-1030 | -8.17689 |
| G721-0221 | -6.21732 | G721-0513 | -8.76759 | G857-1031 | -5.24991 |
| G721-0222 | -6.90937 | G721-0514 | -8.06527 | G857-1032 | -5.45089 |
| G721-0222 | -5.45526 | G721-0517 | -5.72477 | G857-1033 | -6.76847 |
| G721-0223 | -6.25416 | G721-0518 | -7.2347 | G857-1034 | -5.40009 |
| G721-0224 | -7.15457 | G721-0518 | -2.65082 | G857-1035 | -5.33039 |
| G721-0225 | -7.85505 | G721-0519 | -4.86169 | G857-1036 | -5.99136 |
| G721-0225 | -6.03498 | G721-0519 | -5.39654 | G857-1037 | -9.06192 |
| G721-0226 | -8.18708 | G721-0520 | -5.17236 | G857-1038 | -6.05911 |
| G721-0226 | -7.3854 | G721-0521 | -6.49854 | G857-1039 | -7.11901 |
| G721-0227 | -6.16416 | G721-0522 | -6.32719 | G857-1040 | -4.10532 |
| G721-0228 | -4.73519 | G721-0522 | -5.98817 | G857-1041 | -4.95233 |
| G721-0229 | -6.41816 | G721-0523 | -5.57165 | G857-1042 | -5.67319 |
| G721-0232 | -6.09829 | G721-0523 | -4.3146 | G857-1043 | -8.10071 |
| G721-0232 | -4.36795 | G721-0524 | -5.50034 | G857-1044 | -6.6687 |
| G721-0233 | -5.56693 | G721-0525 | -8.07818 | G857-1045 | -3.82252 |
| G721-0526 | -7.07835 | G857-1063 | -8.23754 | G857-1050 | -7.62612 |
| G721-0527 | -5.15455 | G857-1064 | -7.30174 | G857-1050 | -5.36394 |
| G721-0532 | -6.273 | G857-1065 | -4.99849 | G857-1051 | -6.4094 |
| G857-1046 | -7.01045 | G857-1066 | -4.26252 | G857-1052 | -8.96461 |
| G857-1047 | -5.773 | G857-1068 | -4.14665 | G857-1053 | -7.91446 |
| G857-1048 | -5.91752 | G857-1069 | -4.93112 | G857-1054 | -4.98361 |
| G857-1049 | -7.68854 | G857-1070 | -6.38832 | G857-1055 | -6.72249 |
| G857-1050 | -5.48068 | G857-1071 | -7.96143 | G857-1056 | -6.52542 |
| G857-1050 | -5.37639 | G857-1072 | -5.79806 | G857-1057 | -5.57335 |
| G857-1059 | -6.96094 | G857-1061 | -6.91818 | G857-1058 | -4.92063 |
| G857-1060 | -2.31944 | G857-1062 | -6.61382 | G857-1058 | -4.76547 |
